# Contribution of noncanonical antigens to virulence and adaptive immunity in human infection with enterotoxigenic *E. coli*

**DOI:** 10.1101/2020.10.20.348110

**Authors:** FM Kuhlmann, RO Laine, S Afrin, R Nakajima, M Akhtar, T Vickers, K Parker, NN Nizam, V Grigura, CW Goss, PL Felgner, DA Rasko, F Qadri, JM Fleckenstein

## Abstract

Enterotoxigenic *E. coli* (ETEC) contribute significantly to the substantial burden of infectious diarrhea among children living in low and middle income countries. In the absence of a vaccine for ETEC, children succumb to acute dehydration as well as non-diarrheal sequelae related to these infections including malnutrition. The considerable diversity of ETEC genomes has complicated canonical vaccine development approaches focused on a subset of antigens known as colonization factors (CFs). To identify additional conserved immunogens, we mined genomic sequences of 89 ETEC isolates, bioinformatically selected potential surface-exposed pathovar-specific antigens conserved in more than 40% of the genomes (n=118), and assembled the representative proteins onto microarrays, complemented with known or putative colonization factor subunit molecules (n=52), and toxin subunits to interrogate samples from individuals with acute symptomatic ETEC infections. Surprisingly, in this open-aperture approach, we found that immune responses were largely constrained to a small number of antigens including individual colonization factor antigens and EtpA, an extracellular adhesin. In a Bangladeshi cohort of naturally infected children < 2 years of age, both EtpA and a second noncanonical antigen, EatA, elicited significant serologic responses that were associated with protection from symptomatic illness. In addition, children infected with ETEC isolates bearing either *etpA or eatA* genes were significantly more likely to develop symptomatic disease. These studies support a role for more recently discovered noncanonical antigens in virulence and the development of adaptive immune responses during ETEC infections, findings that may inform vaccine design efforts to complement existing approaches.

## Introduction

Enterotoxigenic *Escherichia coli* (ETEC) are one of the commonest causes of childhood diarrhea, accounting for 100s of millions of cases annually (1). This high burden of disease contributes a substantial risk of increased childhood morbidity and mortality (2), (3, 4). Repeated diarrheal infections, including those caused by ETEC, lead to the development of growth stunting and environmental enteropathy, which are life-long consequences of these enteric infections (5). Therefore, preventative efforts, including vaccination, could have a tremendous impact on global health (6). Despite the lack of a licensed ETEC vaccine, two important lines of evidence suggest ETEC vaccine development is feasible. First, controlled human infection models (CHIM) demonstrate that protective immunity develops following ETEC challenge (7, 8). In addition, the frequency of symptomatic infections in young children living in endemic regions wanes substantially with age (9, 10), suggesting that natural infections afford subsequent protection.

ETEC biology, and the extraordinary genetic plasticity of *E. coli*, has complicated the development of a broadly protective vaccine. Canonical approaches have focused primarily on surface features known as colonization factors (CFs) or CS antigens. However, the structural and antigenic diversity of these targets has proved challenging (11). Although toxoids that can elicit neutralizing antibodies against the heat-labile (LT) (12) and heat-stabile toxins (ST) (13) that define the ETEC pathovar are currently under development (14, 15), it is not yet clear whether these alone will afford sufficient, long-lasting protection.

While the ETEC pathovar exhibits high genetic diversity, the recent availability of multiple, genomic sequences from globally diverse ETEC affords the ability to apply reverse vaccinology approaches to the identification of conserved, surface-expressed antigens (16, 17). In addition, microarray-based profiling of immune responses in human volunteers to ETEC challenge has recently highlighted non-canonical antigens recognized during controlled experimental infection.(7, 18).

Application of these approaches to antigen discovery has reinforced the importance of several surface-expressed molecules common to the ETEC pathovar that are not currently targeted in classical vaccine approaches(19). These include two novel secreted molecules, the EtpA adhesin(20) and the EatA(21) autotransporter both originally identified in H10407, an ETEC strain isolated from a case of severe cholera-like diarrhea in Bangladesh. Recent work demonstrates that both antigens are globally distributed in the ETEC pathovar and are more highly conserved than the most common CFs(19, 22). Moreover, they are protective in murine models of infection(23–26) and immunogenic in human challenge trials(7, 18), suggesting that these molecules could provide additional antigenic targets for vaccine development. While much is known about EatA and EtpA under experimental conditions, less is known about their respective roles in natural infections. The present studies were designed to explore the role of these and other potential noncanonical antigens in shaping the adaptive immune response to ETEC infection and to examine their contribution to virulence.

## Methods

### clinical samples used in this study

Specimens used in these studies were obtained from archived studies on ETEC birth cohort carried out in Mirpur in Dhaka city (10) as well as other studies (27) The International Centre for Diarrhoeal Disease Research, Bangladesh (<u>icddr,b</u>). Frozen ETEC isolates were retrieved from storage (−80°C) and duplicate vials were shipped to Washington University and subsequent antigen detection.

### microbial genome analysis and bioinformatic antigen selection

Genomes from 89 clinical ETEC isolates previously collected at icddrb were used to identify conserved surface proteins. Sequence data for all eighty-nine clinical isolates examined in this study are available in GenBank (28). Paired-end Illumina sequence data from each isolate were generated *de novo* and contigs were binned using a previously described protocol (28). The ETEC genomes were compared using LS-BSR as previously described (29–31). The predicted protein-encoding genes of each genome that had ≥90% nucleotide identity to each other were assigned to gene clusters using uclust (32). Representative sequences of each gene cluster were then compared to each genome using TBLASTN (33) with composition-based adjustment turned off, and the TBLASTN scores were used to generate a BSR value indicating the detection of each gene cluster in each of the genomes analyzed. The BSR value was determined by dividing the score of a gene compared to a genome by the score of the gene compared to its own sequence. The predicted protein function of each gene cluster was determined using an ergatis-based (34) in-house annotation pipeline (35). A total of 13,835 non-redundant putative genes (referred to here as ‘centroids’) were extracted from the eighty-nine genomes.

All 13,835 centroids in this study were subjected to a reverse vaccinology pipeline (Institute for Genome Sciences, Maryland, USA) to identify those features that contained features that suggested they were surface exposed. An additional subtractive analysis was conducted by filtering centroids (BLASTx and BLASTn) against the genome contents of six *E. coli* commensal and laboratory strains, yielding 6,444 ETEC pathovar-specific centroids. These data were further refined by selecting centroids with a blast score ratio (BSR)(36) ≥ 0.8 (i.e. highly conserved) and present in at least 40% of the clinical isolates, yielding 316 conserved, virulence-linked genetic features for further analysis. BLASTx was next used to assign a putative function to these virulence-linked centroids. This analysis was coupled with results from pSORTv3.0 (37), SubLoc (38), and CELLO (39) to predict subcellular localization, altogether resulting in the down-selection to 118 potential surface-expressed molecules These features were complemented with all known and putative colonization factor subunits (n=52), toxin subunits, and subdomains of novel antigens for inclusion on the microarrays (supplemental dataset 1).

### microarray production

Antigen-encoding regions selected for the microarrays were amplified by PCR using primers listed in supplemental dataset 2, and constructed as previously described(7, 18, 40) Recombinant versions of select antigens including EtpA, EatA, LT-A, LT-B, YghJ, ST-H, and EaeH were also included on the arrays.

### microarray processing

Microarrays were shipped to icddrb where they were rehydrated for 10 minutes with 100 ul Array Blocking Buffer. *E. coli* lysate was reconstituted in a final volume of 20% in blocking buffer. Antibody in Lymphocyte Supernatant (ALS) prepared from blood of ETEC patients were diluted 1:2.5 in the resuspended lysate followed by loading onto the microarrays and incubated in the dark for 2 hours at 25°C on a rotating platform. Microarrays were then washed 3x with TBS-T (0.05% Tween in TBS, pH 7.5) followed by incubation for 5 minuites in TBS-T at 25°C. This process was repeated once with TBS followed by a final wash in distilled water. Slides were dried by centrifugation (10 minutes at 500 x *g*) then stored in desiccated boxes prior to shipping to the Felgner Laboratory, University of California, Irvine.

### non-canonical antigen ELISA

384 well plates (<u>Corning, product number 3540</u>) were coated with recombinant EatA passenger domain (rEatp, 10 micrograms/milliliter in carbonate buffer [15 mM Na_2_CO_3_, 35 mM NaHCO_3_, 0.2g/l NaN_3_, pH 9.6]) or recombinant EtpA (rEtpA, 1 microgram/milliliter in carbonate buffer) and shipped to icddr,b, being maintained at 4°C prior to use. The ELISA plates were manually washed three times with PBS-T (PBS with 0.05% tween) including brief centrifugation for 30 seconds at 200 x *g* on a tabletop centrifuge between washes. Plates were rehydrated with 1% BSA in PBS-T overnight at 4°C. The following day, serum or plasma samples and plates were warmed to ambient temperature (~25°C), serum was diluted 1: 200 in PBS-T with 1%BSA and briefly vortexed. 10 μl of diluted serum was added to the plates, centrifuged as above, sealed, and incubated at 37°C for 1 hour. After incubation, plates were washed 3 times with PBS-T as described above. 10 μl of HRP-conjugated anti-human IgG (Jackson ImmunoResearch Laboratories, <u>Cat 309-035-006</u>, West Grove, PA) was diluted 1:2000 in 1% BSA in PBS-T followed by incubation and washing as above. ELISA plates were read using 10 ul of 3,3’,5,5’-tetramethylbenzidine (TMB) substrate (Seracare, <u>Cat# 50-76-00</u>, Milford, MA) and the Vmax of the reaction was determined using a BioTek Plate reader with Gen5 Take3 software (v.2.00.18). Due to variations between ELISA plates, we analyzed data independently for each plate and in combination, adjusting for age to control for repeated measures.

### strain characterization by PCR and immunoblotting

Frozen glycerol stocks of ETEC strains maintained at −80°C were used to inoculate lysogeny broth (LB) for overnight growth at 37°C, 250 rpm. 1 μl of the overnight culture was diluted in 100 μl of PBS of which 1 μl was used as the DNA template in initial PCR screening with primers in supplementary table 1. The thermocycler conditions for *eatA* and *etpA* were denaturation for 5 minutes at 95°C with 30 amplification cycles utilizing 95°C for 30 seconds, 52°C for 30 seconds, and 72°C for 2 minutes. The toxin multiplex assay (genes *eltB*, *estH*, and *estP*) were conducted as follows; 5 minutes at 95°C with 32 cycles of amplification using 94°C for 15 seconds, 55°C for 15 seconds, and finally 72°C for 30 seconds. Amplicons were visualized as before using a 0.8% agarose gel with ethidium bromide. The H10407 strain (*eatA*, *etpA*, *estH*, *estP*, and *eltB* positive) was used as a positive control in for assays. To adjudicate discordant results PCR was performed using gDNA extraction with the Invitrogen PureLink Quick Plasmid Miniprep Kit (<u>Cat# K210010</u>, Thermo Fisher, Waltham, MA) Miniprep kit. If toxin multiplex PCRs were negative, isolates were deemed to have lost their original plasmid during storage, transportation, or culture passage and subsequently excluded from analysis. Immunoblotting for EatA and EtpA were performed on TCA-precipitated culture supernatants as previously described (19) using affinity-purified polyclonal rabbit antibodies against the passenger domain of EatA (21) or EtpA (20) (dilutions 1:1,000 and 1:5,000, respectively) Primary antibodies were detected using Horseradish Peroxidase (HRP)-conjugated anti-rabbit IgG secondary antibody (1:5,000 dilution, Invitrogen #A16110) for 1 hour at room temperature. HRP was detected with ECL Western blotting substrate (Bio-Rad, #ABIN412579).

### statistical analysis

Categorical outcomes were analyzed using chi-square tests, Fisher’s exact tests, or age-adjusted logistic regression analyses as appropriate. Serum data were analyzed using a linear repeated measures model with a compound symmetry covariance structure. *p*-values < 0.05 were considered significant. Analyses were conducted using SAS version 9.4 (SAS Institute Inc., Cary, NC, USA) or SPSS v.24 (IBM, Armonk, NY, USA), or GraphPad Prism v9.0.0.

### Ethics Statement

These studies were approved by the Research Review and Ethical Review Committee of (icddr,b) and the Institutional Review Board of Washington University School of Medicine in Saint Louis.

## Results

### Antibodies following natural infection recognize a finite repertoire of ETEC proteins

Both human experimental models(7) as well as natural infections(10) demonstrate that prior infection with ETEC affords substantial protection against symptomatic disease. Elucidation of the nature of protective adaptive immune responses to these mucosal pathogens can therefore inform vaccine development. While the majority of earlier ETEC vaccinology efforts have centered on colonization factor antigens, the present studies were designed to broadly profile antigenic responses and to assess the role of recently characterized non-canonical antigens. To assess the breadth of immune responses to ETEC during acute natural infection, we designed protein microarrays containing all known and putative colonization factor antigen subunits, and additional predicted surface-expressed proteins conserved in more than 40% of the ETEC pathovar including EtpA and EatA, secreted antigens expressed by a majority of ETEC strains in a global collection of isolates(19).

Despite the inclusion of multiple candidate surface molecules on the array predicted to be conserved among strains in Bangladesh from our *in silico* analysis, immune responses following infection were largely constrained to a small group of antigens including EtpA and EatA (figure 1A), LT (supplementary figure 1), select colonization factor subunits (supplementary figure 2) including CssB, one of two components of the CS6 polymer(41), a predominant immunogenic antigen among strains circulating in Bangladesh(27). Compared to control specimens obtained outside of the ETEC endemic area, both EatA and EtpA exhibited high levels of reactivity. Notably, for patients infected with EtpA-expressing strains, EtpA responses were significantly higher at day 30 following infection than those observed immediately following admission, whereas the converse was true in patients admitted with EtpA-negative strains. (figure 1B)

**Figure 1.**
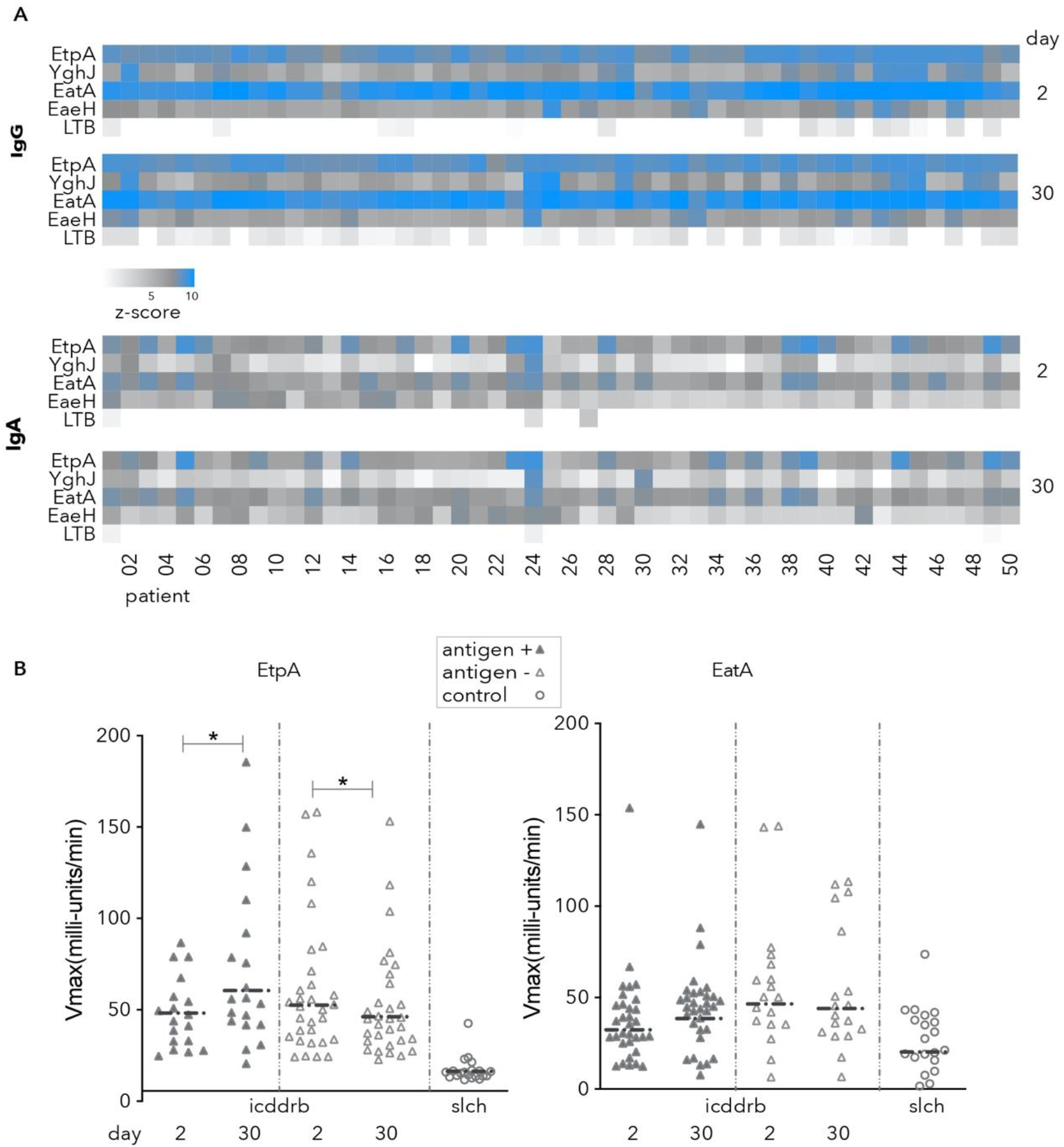
serologic response to non-canonical antigens following natural infection. **A.** heatmap indicates log 2 transformed z-score data indicating ETEC protein microarray responses from day 2, and 30 following presentation to icddrb to four non-canonical antigens EtpA, YghJ, the passenger domain of EatA, EaeH; and the B-subunit of ETEC heat-labile toxin (LT-B). **B.** kinetic ELISA responses to EtpA and EatA following infection. Data are segregated by the presence (closed symbols), or absence (open symbols) of each respective antigen in the strain recovered at presentation. Negative control samples from Saint Louis Children’s Hospital (slch) are shown as open circles. #<0.05 by Wilcoxon matched-pairs signed rank test.

In an open-aperture assessment of ALS specimens (42, 43) obtained from adults hospitalized at icddr,b Hospital in Dhaka, Bangladesh or from patients recruited at the Mirpur field site with acute symptomatic diarrheal illness, we again noted that immune responses following infection were largely constrained to a relatively small group of antigens including CS6, EtpA and EatA (supplementary dataset 1). When parsing antigen profiles of the infecting strain, we found that those infected with EtpA-expressing ETEC exhibited significant increases in both ALS IgA (*p*=0.005), and IgG (*p*=0.02) responses in the week following infection relative to those infected with EtpA-negative strains (figure 2). As anticipated, we also observed significant increases in ALS immunoreactivity to the CssB subunit of CS6 that correlated with the production of CS6 by the infecting strain (supplementary figure 3).

**Figure 2.**
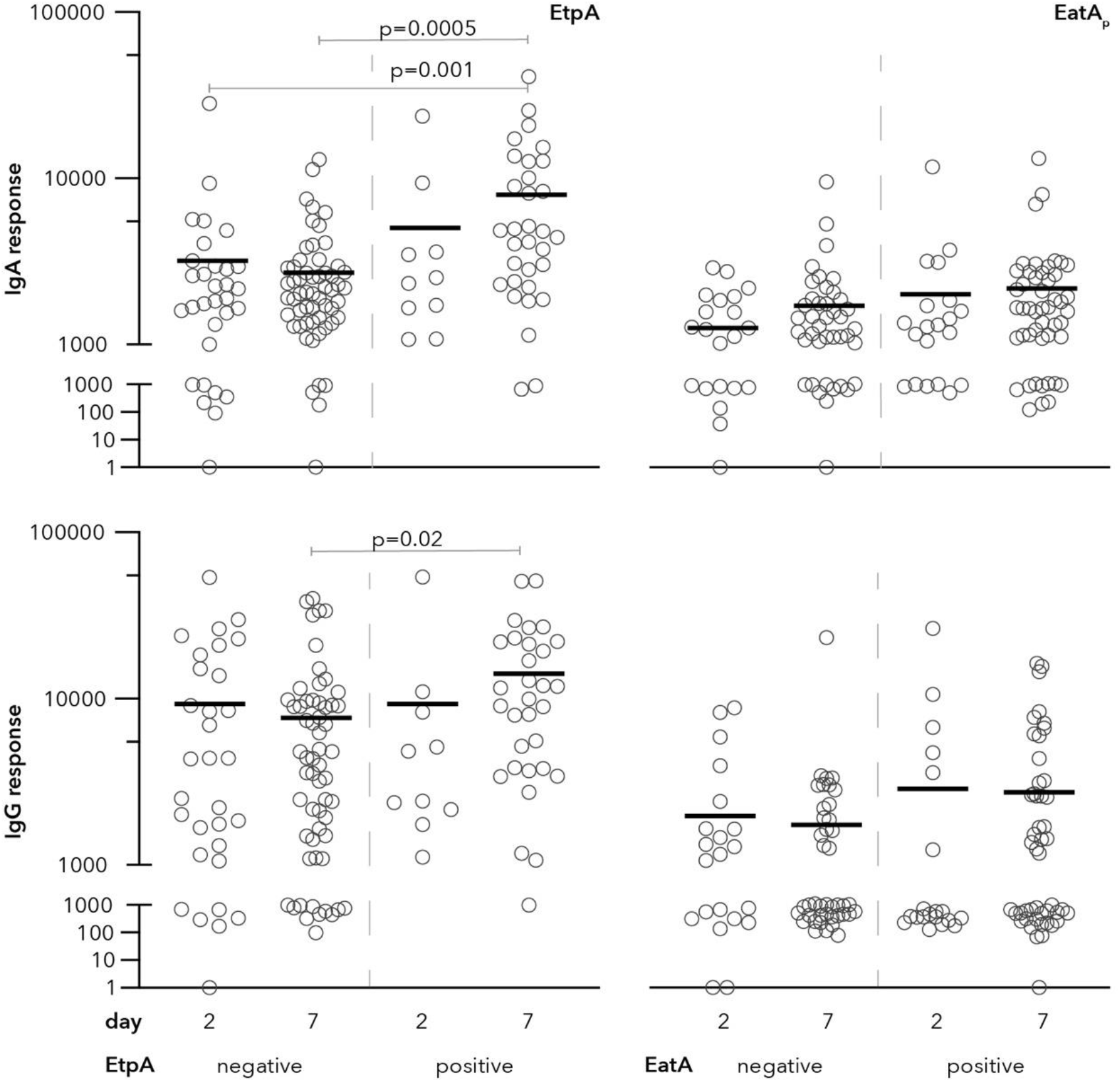
ALS responses to EtpA or EatA. Shown are microarray data for IgA (top panels) and IgG responses to EtpA (left) and the passenger domain of EatA (EatAp, right) on days 2, and 7 following hospitalization. Data in each graph are segregated according to antigen expression in the infecting strain (negative or positive). p values reflect Kruskall-Wallis, with post-hoc analysis using Dunn’s test adjusted for multiple comparisons for between group analysis.

### EatA and EtpA are immunogenic in young children

Data from recent CHIM studies(7, 18) as well as earlier data from patients with natural ETEC infections(22), indicate that adults develop robust immune responses to non-canonical antigens including EtpA and EatA. However, in endemic areas young children are the population most severely impacted by ETEC with incidence declining after 24 months of age, presumably as protection develops subsequent to infection. Therefore, we examined sera from a cohort of Bangladeshi children followed from birth through 2 years of age (10) to profile development of antibody responses to EatA and EtpA over time (figure 3). During the first month of life, the majority of children were observed to have elevated IgG responses to both EatA and EtpA, presumably reflecting passive transfer of maternal antibodies(44). As anticipated, responses to both antigens decreased by three months of age, while mean responses to each antigen increased significantly through 24 months of age, likely reflecting early childhood infections with strains expressing EtpA and EatA.

**Figure 3.**
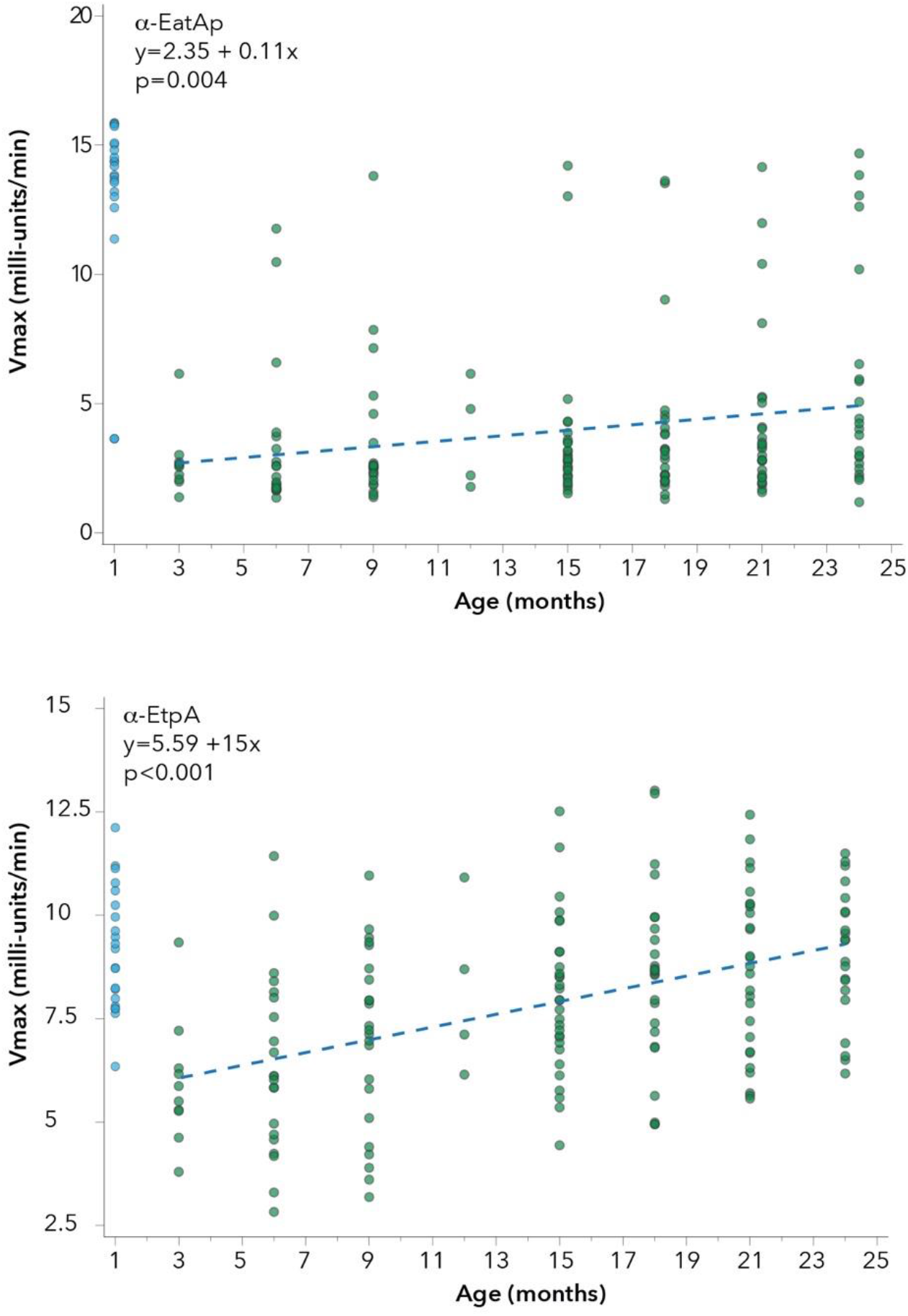
Anti-EtpA or Anti-EatA IgG responses increase with age. Shown are representative kinetic ELISA data for serum IgG samples obtained from children ages 1-24 months enrolled in a birth cohort study. Scatter plots of Anti-EtpA and Anti-EatA IgG plotted against data for ages 3-24 months with regression lines from linear repeated measures models overlaid (dotted line) demonstrate significant increases over time in responses to the passenger domain of EatA (EatA_p_, top) and EtpA (bottom). See Figure S4 for additional plots.

### Anti-EtpA or EatA responses relative to symptomatic diarrhea

Immunologic correlates of protection against ETEC are currently unknown(45). The majority of clinical studies to date have examined the impact of prior infection with strains producing particular colonization factors and/or LT(9) as well as antibody acquisition on subsequent risk of infection with similar strains(46, 47). We hypothesized that because EtpA and EatA are relatively common antigens in the ETEC pathovar(19), higher antibody responses to these antigens may be associated with subsequent protection against symptomatic infection. After excluding antibody responses at one month of age, we examined the IgG antibody responses to EtpA and EatA preceding detection of ETEC in either symptomatic or asymptomatic children between four and twenty-four months of age. Interestingly, we observed elevated responses to both antigens prior to detection of ETEC in asymptomatic children detection relative to symptomatic cases (figure 4), perhaps reflecting the overall mitigating impact of prior exposure on development of diarrheal illness.

**Figure 4.**
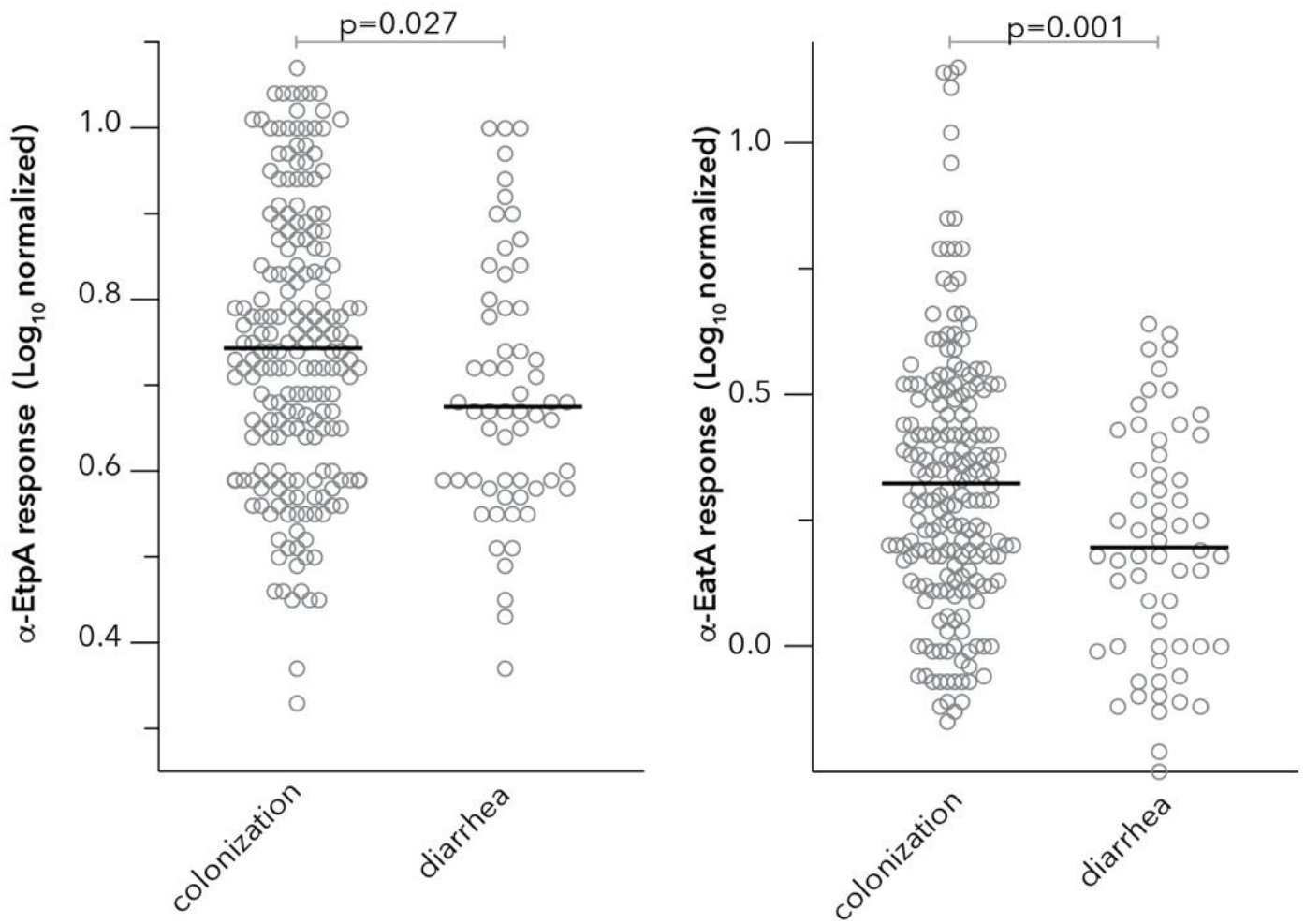
Serum IgG responses preceding asymptomatic ETEC colonization and diarrhea. Shown are peak serum IgG responses for EtpA (left) or the EatA passenger domain (right) preceding either asymptomatic colonization or diarrheal illness with ETEC. Data shown are Log_10_ transformed IgG antibody responses determined by kinetic ELISA. Bars represent mean values.

### etpA relative to blood group A upon first ETEC exposure

Recent studies have shown that the EtpA adhesin engages host cells via lectin interactions with N-acetylgalactosamine residues, preferentially when presented as the terminal sugar on blood group A glycans displayed on intestinal epithelia. Also, when challenged with ETEC H10407, an EtpA-producing strain, human volunteers of A blood group are more likely to exhibit severe symptomatic illness(48), recapitulating the observation that symptomatic ETEC infections were more common in children with blood type A or AB in birth cohort studies of Bangladeshi children(10). We therefore examined this cohort for potential associations between *etpA*, blood type, and disease status during natural ETEC infections. In limiting analysis to the first ETEC isolation to avoid confounding effects of repeated exposures, 41.2% of children that were blood type A or AB had symptomatic diarrhea during their first ETEC infection (table 1) compared to 30.6% of those that were blood group B or O (30.6%, *p* = 0.187). Blood group A or AB individuals were somewhat more likely to have an *etpA* positive strain recovered from their first infection (74.5% vs. 66.7%, 0R = 1.462, 95%CI 0.693-3.082), although these data were not statistically significant.

**table 1.** associations of blood group, etpA expression and diarrheal disease.

|  |  |  |  |  |
| --- | --- | --- | --- | --- |
| table 2. association of blood group, *etpA*, and diarrheal illness |  |  |  |  |
|  | clinical presentation |  | *etpA* status |  |
|  | asymptomatic | symptomatic | negative | positive |
| blood type | n (%) | n (%) | n (%) | n (%) |
| O or B | 75 (69.4) | 33 (30.6) | 36 (33.3) | 72 (66.7) |
| A or AB | 30 (58.8) | 21 (41.2) | 13 (25.5) | 38 (74.5) |
|  |  | p=0.187* |  | p=0.317<sup>†</sup> |
| * comparison of symptomatic infections in A/AB vs B/O |  |  |  |  |
| <sup>†</sup> comparison of *etpA* positivity in A/AB vs B/O individuals with first episodes of diarrhea |  |  |  |  |
| p values determined by chi-square test |  |  |  |  |

### association of *eatA* and *etpA* with virulence

Although both EatA, a mucin-degrading serine protease, and the EtpA blood group A lectin are secreted by a diverse population of ETEC strains(19), and contribute to virulence phenotypes *in vitro* as well as in small animal models of ETEC infection(23, 26, 49, 50), the role played by these antigens in human infections has yet to be explored in detail. To explore the association of *eatA* and *etpA* with symptomatic ETEC infection, we examined isolates collected in a birth cohort study in which stools were collected at monthly intervals in asymptomatic children (asymptomatic colonization) of during surveillance for diarrhea (symptomatic infection) (10). Notably, the presence of *etpA* or *eatA* significantly increased the odds of having symptomatic diarrhea (unadjusted odd ratios of 2.1 and 3.1, respectively (table 2). Similarly, after adjusting for age we observed significant associations between the presence of either EtpA (adjusted odds ratio 1.98, *p*=0.007) or EatA (adjusted odds ratio 2.91, *p*<0.001) and development of diarrheal disease.

**table 2.**
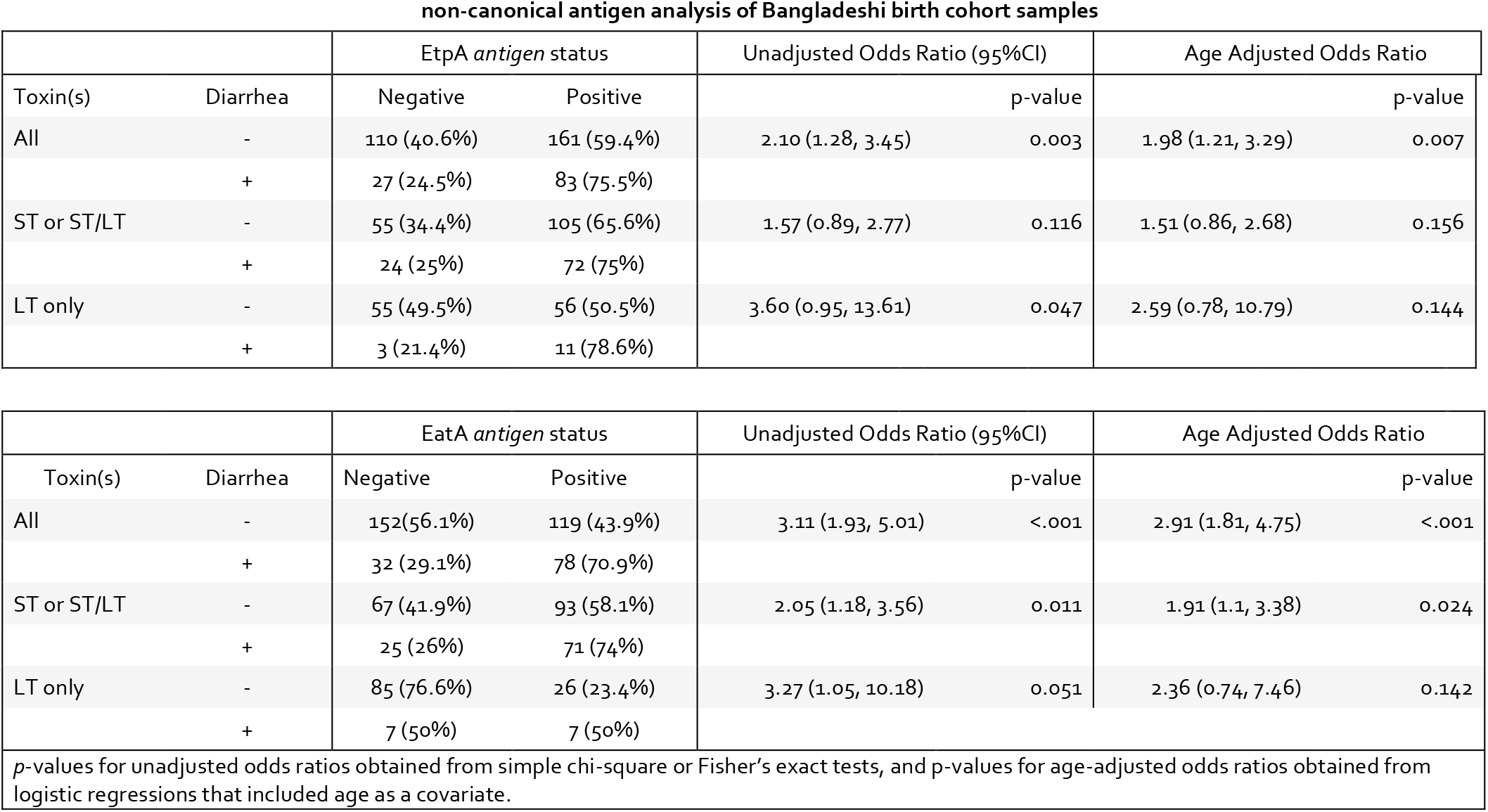
relationship of *eatA*, and *etpA*, to symptomatic ETEC.

non-canonical antigen analysis of Bangladeshi birth cohort samples
|  |  | EtpA *antigen* status |  | Unadjusted Odds Ratio (95%CI) |  | Age Adjusted Odds Ratio |  |
| --- | --- | --- | --- | --- | --- | --- | --- |
| Toxin(s) | Diarrhea | Negative | Positive | p-value |  | p-value |  |
| All | - | 110 (40.6%) | 161 (59.4%) | 2.10 (1.28, 3.45) | 0.003 | 1.98 (1.21, 3.29) | 0.007 |
|  | + | 27 (24.5%) | 83 (75.5%) |  |  |  |  |
| ST or ST/LT | - | 55 (34.4%) | 105 (65.6%) | 1.57 (0.89, 2.77) | 0.116 | 1.51 (0.86, 2.68) | 0.156 |
|  | + | 24 (25%) | 72 (75%) |  |  |  |  |
| LT only | - | 55 (49.5%) | 56 (50.5%) | 3.60 (0.95, 13.61) | 0.047 | 2.59 (0.78, 10.79) | 0.144 |
|  | + | 3 (21.4%) | 11 (78.6%) |  |  |  |  |

table 2. relationship of *eatA*, and *etpA*, to symptomatic ETEC.
|  |  | EatA *antigen* status |  | Unadjusted Odds Ratio (95%CI) |  | Age Adjusted Odds Ratio |  |
| --- | --- | --- | --- | --- | --- | --- | --- |
| Toxin(s) | Diarrhea | Negative | Positive | p-value |  | p-value |  |
| All | - | 152(56.1%) | 119 (43.9%) | 3.11 (1.93, 5.01) | <.001 | 2.91 (1.81, 4.75) | <.001 |
|  | + | 32 (29.1%) | 78 (70.9%) |  |  |  |  |
| ST or ST/LT | - | 67 (41.9%) | 93 (58.1%) | 2.05 (1.18, 3.56) | 0.011 | 1.91 (1.1, 3.38) | 0.024 |
|  | + | 25 (26%) | 71 (74%) |  |  |  |  |
| LT only | - | 85 (76.6%) | 26 (23.4%) | 3.27 (1.05, 10.18) | 0.051 | 2.36 (0.74, 7.46) | 0.142 |
|  | + | 7 (50%) | 7 (50%) |  |  |  |  |

p-values for unadjusted odds ratios obtained from simple chi-square or Fisher's exact tests, and p-values for age-adjusted odds ratios obtained from logistic regressions that included age as a covariate.

The *eatA* gene (21) and *etpBAC* locus(20) encoding the two-partner secretion system responsible for EtpA secretion, were originally identified on the p948 plasmid of ETEC strain H10407, which also encodes the gene for STh (51), and our earlier studies suggested that both loci are more commonly associated with ST-producing strains (19). Importantly, large epidemiological studies have demonstrated an association between ST or ST/LT-producing ETEC and more severe disease relative to LT-only producing ETEC (52, 53). Similarly, we again found an association between ST-producing ETEC and symptomatic diarrhea, where 59.0% of colonizing ETEC isolates encode STh or STp (*estH* or *estP* positive) compared to 87.3% of diarrhea-associated isolates (adjusted odds ratio 4.66 [95% CI, 2.62, 8.85, *p* < 0.001]). We therefore asked whether the *eatA* or *etpA* associations with virulence were independent of ST. The presence of either gene was associated with higher risk of diarrheal illness independent of ST, although only the presence of *eatA* was significantly associated with illness adjusted for age. Collectively, however these data suggest that these more recently discovered non-canonical antigens, now frequently referred to as “accessory” virulence factors, could be important contributors to ETEC disease.

## Discussion

ETEC were initially discovered in patients presenting with severe diarrheal illness that mimicked clinical cholera (54–56). Following seminal discoveries of the heat-labile (LT) and heat-stable (ST) toxins that define ETEC, and initial characterization of plasmid-encoded colonization factor antigens (CFs), a canonical approach to vaccine development focused on LT and CFs emerged. However, subsequent studies have revealed that the molecular pathogenesis of ETEC likely involves a number of other plasmid as well as chromosomally encoded features that may potentially expand the repertoire of target “non-canonical” antigens for use in ETEC vaccine development.

Among antigens that are unique to the ETEC pathovar are two high molecular weight secreted proteins, EtpA and EatA. The relative conservation of genes encoding their corresponding secretion systems within the ETEC pathovar(18, 19), their immunogenicity during natural(57) and experimental human infection(7, 18), and contribution to virulence *in vitro* and small animal studies have highlighted their potential utility as vaccine candidates. Nevertheless, our understanding of the importance of these antigens to ETEC virulence continues to evolve. Recent studies have revealed that the secreted 110 kD passenger domain of the EatA autotransporter protein functions as a mucin-degrading enzyme, capable of dissolving the MUC2 matrix that covers the surface of enterocytes, the target for ETEC binding and toxin delivery(23). EtpA, secreted by two-partner secretion mechanism that requires both the EtpB outer membrane pore and EtpC, a glycosyltransferase(20), functions as an adhesin by bridging the bacteria(50) and GalNAc-containing host cell glycans present on enterocytes(48). However, despite an emerging understanding of the function of these molecules, very little is known about their contribution to disease in human hosts.

The present studies extend earlier observations to a cohort of naturally infected children in Bangladesh(10) and suggest that these non-canonical antigens play critical roles in determining the outcome of ETEC infections. The finding that genes encoding these antigens are significantly associated with the development of symptomatic infection may have important implications for the interpretation of large-scale epidemiologic studies that have employed population attributable fraction methodology in which ETEC detected in cases of diarrheal illness are compared to asymptomatically colonized controls(58). The present studies would seem to suggest that additional characterization of ETEC beyond the pathovar-defining heat-labile or heat-stabile toxins could be required to accurately assess the contribution of ETEC to the global burden of diarrheal disease.

In general, expanded open-aperture assessment of immune responses to natural ETEC infections appears to reaffirm earlier observations in human volunteer studies(7). Namely, that there are relatively few immunogenic targets in the potential repertoire of ETEC surface molecules, with EtpA and EatA predominating among the pathovar-specific antigens.

Although we observed higher IgG serum antibody responses to both EtpA and EatA in children who were simply colonized with ETEC compared to those with diarrhea, suggesting that these antigens could afford some protection against symptomatic illness, these findings need to be interpreted cautiously. Both EtpA and EatA are relatively common antigens among strains circulating in Bangladesh, therefore the identification of antibodies could simply reflect prior infection that mitigates infection through responses to other antigens. In addition, correlates of protection for ETEC, as well as the protective role of serum IgG in enteric infections remain unclear with mucosal IgA responses considered key to protection (45).

Altogether, however the findings reported here suggest that antigens which have not been part of traditional approaches to vaccine development may play important roles in virulence, and in acquired immunity to ETEC. Further studies will clearly be needed to examine the efficacy of these more recently discovered antigens as protective immunogens.

## Acknowledgements

These studies were supported in part by funding from National Institute of Allergy and Infectious Diseases (NIAID) of the National Institutes of Health (NIH) under Award Numbers R01AI089894, R01AI126887 (jmf), K23AI30389 (fmk) and The Department of Veterans Affairs I01BX004825 (jmf). The icddr,b is supported by the Governments of Bangladesh, Canada, Sweden, and the UK. James M. Fleckenstein is listed the inventor on patent 8323668

## supplementary data

### supplementary figures

**supplementary figure 1.**
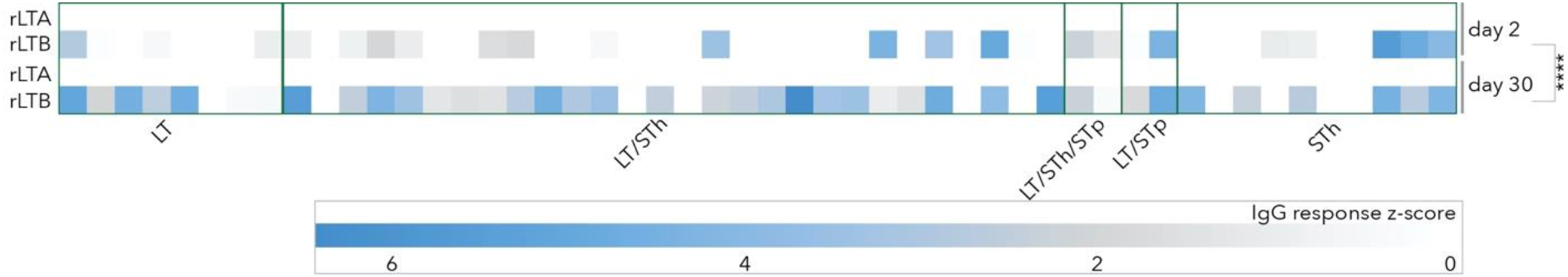
responses to LT subunits LT-A and LT-B following infection. Shown are array z-score data from days 2 and 30 following presentation to icddrb. Data are segregated by the toxin profile of the ETEC strain isolated at presentation. ****=p<0.0001 by Wilcoxon matched pairs comparison of day 2 and day 30 LT-B responses.

**supplementary figure 2.**
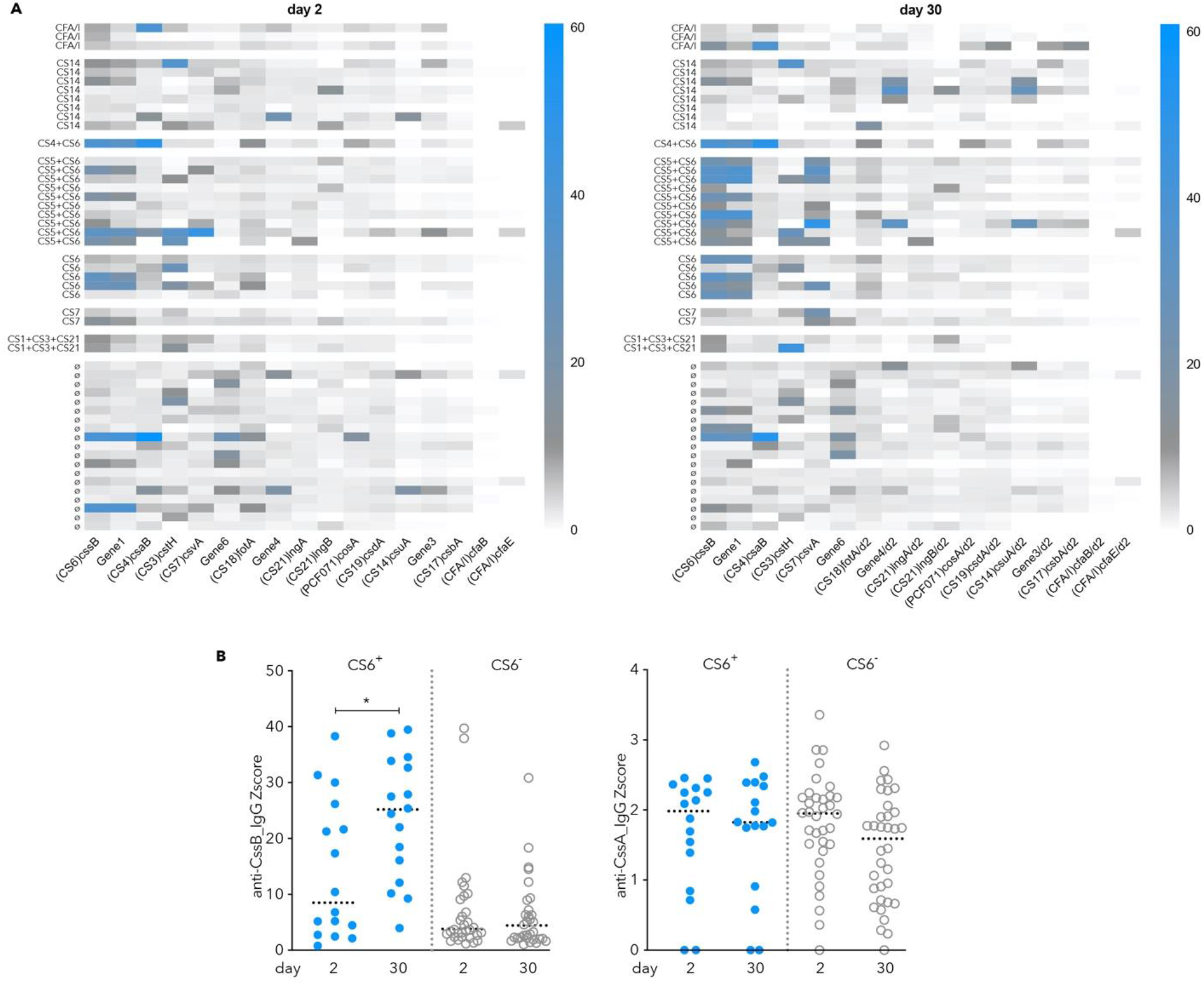
(**A**) Serum IgG responses to colonization factor subunits following ETEC infection, on day 2 and day 30 following presentation to icddrb. Shown are normalized zscore data for select CF antigens (bottom) printed on the microarrays for 50 individuals infected with ETEC. Antigens expressed by the infecting strain for each patient are shown at the left of the heatmap. Responses following infection with CS6-expressing strains are outlined in green. **B**. Serologic responses to CssB (left), and CssA (right) subunits of CS6 following infection. Data are parsed by antigen expression in the infecting strain with symbols in blue indicating CS6+ samples. *p=0.016 by Wilcoxon matched-pairs signed rank testing.

**supplementary figure 3.**
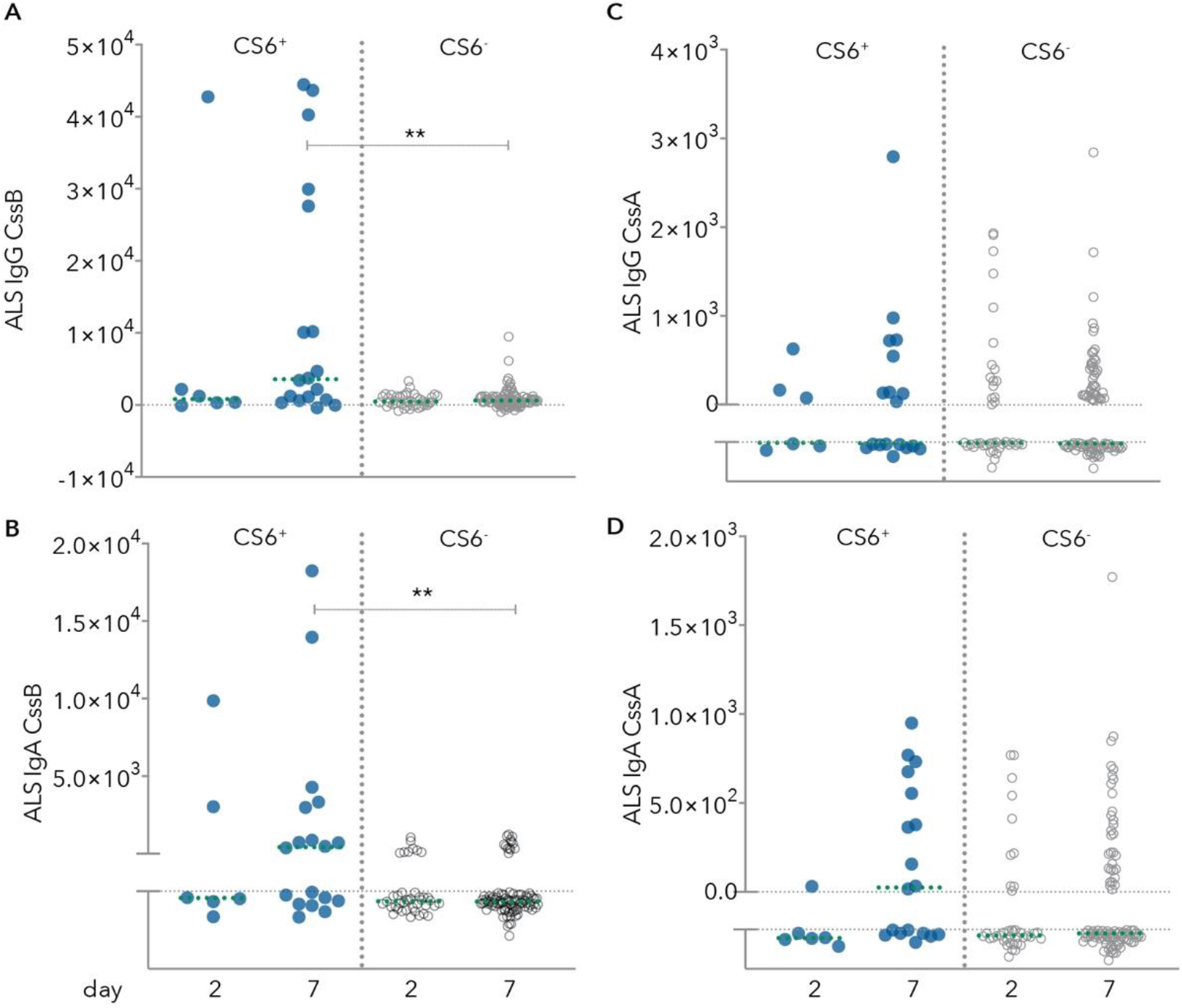
ALS responses to CS6 subunits following ETEC infection. (**A**) IgG response to CssB. (**B**) IgA response to CssB. **p<0.01, Kruskal-Wallis post-hoc analysis using Dunn’s test adjusted for multiple comparisons. (**C**) IgG response to CssA. (**D**) IgA response to CssA.

### supplementary tables

**supplementary table 1.**
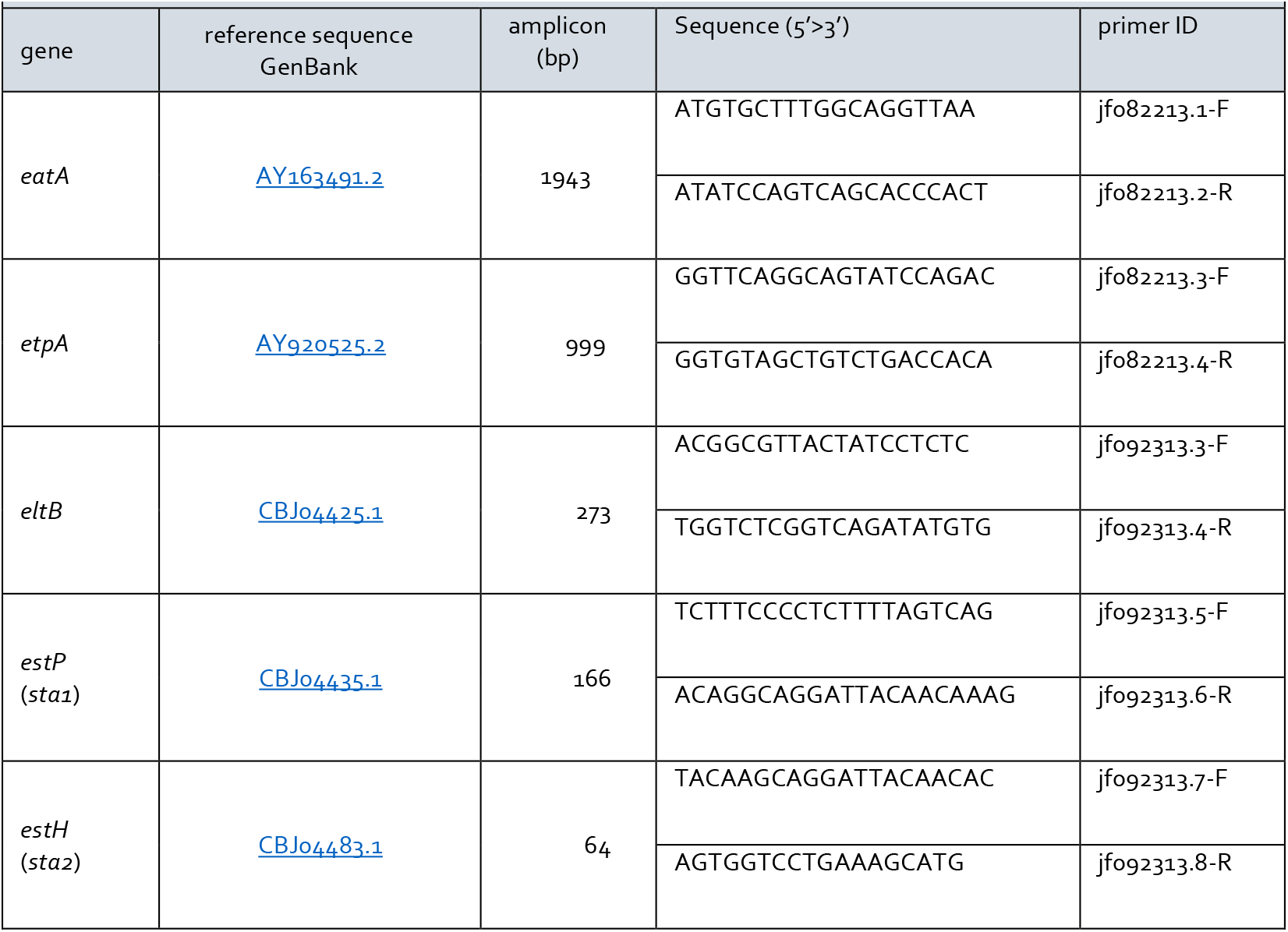
primers used in strain interrogation

